# LuxSit Pro and sPro: Next-Generation Designed Luciferases for Bioluminescent Reporting and Complementation-Based Detection

**DOI:** 10.1101/2025.07.21.665742

**Authors:** Renan Vergara, Jonathan Wagner, Paursa Kamalian, Joseph L. Harman, Nicholas Lara, Elyse S. Fischer, Gonçalo J. L. Bernardes, Alfredo Quijano-Rubio, Daniel-Adriano Silva

## Abstract

Bioluminescent systems are widely used as molecular reporters in life sciences due to their low background, broad dynamic range, and the key advantage of not requiring external excitation. However, natural luciferases often lack the brightness, stability, and long-lasting light emission needed to support a wide range of high-sensitivity applications. Here, we describe our next-generation luciferase solution: LuxSit Pro and its substrate, Luxterazine (LTZ). LuxSit Pro is a de novo luciferase optimized for steady brightness and structural stability through an iterative combination of computational and experimental methods. Composed of only 117 amino acids, LuxSit Pro is currently the smallest highest-performing known luciferase, exhibiting exceptional brightness and biochemical stability with robust recombinant expression in both mammalian and bacterial systems. Notably, LuxSit Pro was deliberately designed devoid of lysine and cysteine residues which enormously expands its potential applications. This property renders the reporter largely resistant to post-translational modifications such as ubiquitination and also enables site-directed chemical modifications, as lysines or cysteines can be reintroduced at specific sites to serve as chemical handles for augmenting or adapting the enzyme’s function. The novel synthetic substrate, Luxterazine (LTZ), is a compound that enables LuxSit Pro to deliver bright, steady, blue light emission (∼490 nm) and offers improved solubility compared to existing Coelenterazine analogs. Finally, we also developed LuxSit sPro, a highly sensitive two-component complementation system for the quantitative detection of molecular interactions. The broad potential applications of LuxSit Pro and sPro, in combination with Luxterazine, are poised to drive the development of next-generation reporters and biosensor systems for the research and *in vitro* diagnostics fields.

## Introduction

Bioluminescent systems have become landmark tools in research and medical applications. To date, researchers have identified over 40 distinct luciferase-luciferin pairs across diverse organisms, with comprehensive structural characterization available for 11 of these systems [1]. Among the most prominent natural luciferases are the ATP-dependent luciferases (e.g., Firefly and Click Beetle) which use D-luciferin as substrate, and the ATP-independent marine luciferases (e.g., Renilla, Gaussia, and Oplophorus gracilirostris) which employ the substrate Coelenterazine (CTZ) [2]. As reporters, luciferases provide important advantages over fluorescent reporters, due to their minimal background and absence of photobleaching (although some luciferases suffer from active site damage due to catalysis). However, natural luciferases can also present significant general limitations as reporters, such as relatively weak and rapidly decaying light emission (due to substrate depletion and/or substrate induced inhibition), high sensitivity to pH changes, poor thermostability, and, in many cases, dependence on disulfide bonds (or even single cysteine residues) to maintain their activity, which limits their use in reducing environments [3].

The Promega NanoLuc luciferase was developed over a decade ago to address many of the limitations of natural luciferases [4]. The NanoLuc–furimazine system is approximately 150 times brighter than Firefly or Renilla luciferases, generates glow-type luminescence, and performs reliably under a wide range of experimental conditions. A few years later, Promega introduced NanoBiT, a split NanoLuc complementation system designed for studying bimolecular interactions [5]; today NanoLuc and the NanoBiT systems are broadly used to study biological systems and to develop biosensors [6]. Despite NanoLuc’s widespread adoption for many applications, no alternative systems have been yet developed that improve over NanoLuc’s properties– until now.

Recently, breakthrough advances in computational protein design allowed David Baker’s group to create the first de novo luciferase enzyme, LuxSit-i [7]. With a molecular weight of approximately 13.5 kDa, and remarkable thermostability (Tm > 90°C), LuxSit-i (part of Monod Bio’s exclusively licensed LuxSit™ portfolio) catalyzes the (ATP-independent) oxidation of the synthetic substrate diphenylterazine (DTZ), with a peak light emission at 480 nm. Recently, Ye’s group expanded the emission of LuxSit-i to a broader color palette through highly-efficient BRET fusions with fluorescent proteins [8]. Despite these advances, LuxSit-i exhibits approximately 300-fold lower brightness than NanoLuc, limiting its use in applications requiring sensitive reporters.

Here we report the development of “LuxSit Pro” an extensively optimized de novo luciferase and its novel soluble substrate Luxterazine (LTZ, a proprietary LuxSit Pro Substrate) to deliver a robust reporter with unique biophysical and biochemical properties not found in any existing luciferases. LuxSit Pro exhibits unparalleled glow-type brightness compared to natural luciferases, and displays exceptional thermal stability (melting point >95°C). Through computational design, we have removed all lysine residues from the protein, addressing the main concerns surrounding the use of native luciferases (or any other fusion protein reporters in general) in targeted protein degradation (TPD) studies [9]. This lysine- and cysteine-free characteristic of LuxSit Pro also provides researchers with a tool for precise and simple site-specific conjugation by reintroducing lysine or cysteine residues at defined positions when needed.

We also developed the two component LuxSit sPro complementation system to enable the study of molecular interactions. LuxSit sPro is composed of two inactive poly-peptides with low-affinity for each other (K_D_ = 90 µM). When forced to be in proximity (by colocalization domains, such as mAbs or chemically induced dimers) the two peptides interact efficiently and reconstitute activity. The two proteins that compose the system are named αLux (103 a.a.) and βit (12 a.a.). LuxSit sPro offers a versatile and sensitive, plug-and-play platform for studying biomolecular interactions (both, in solution or in cells) by using simple genetic fusions, as well they enable the straightforward engineering of sensitive and quantitative homogeneous biosensors for *in vitro* applications. LuxSit sPro retains the key attributes of LuxSit Pro, including compact size, high stability, strong luminescent output, and absence of lysines and cysteines. Upon complementation (by proximity/co-localization) the LuxSit sPro system can achieve full enzymatic activity, comparable to that of its “intact” LuxSit Pro counterpart.

LuxSit Pro and LuxSit sPro provide researchers with a novel bioluminescent toolkit carefully designed to synergize with current technologies to enable the development of next-generation tools; Ultimately Luxit Pro and sPro are primed to kickstart a revolution for the engineering of next-generation biological and biomedical reporter applications.

## Results

### Optimization of the LuxSit Pro - Luxterazine (luciferase-luciferin) pair

The LuxSit Pro bioluminescent system was developed through computational modeling starting from LuxSit-i [7], then subject to iterative rounds of directed evolution in yeast, computational protein redesign, and experimental screening of novel substrates. Yeast surface display was chosen as the empirical experimental system (over bacterial hosts) to ensure full accessibility of the proteins to soluble substrates during the experimental optimization process.

The first round of optimization consisted of a single-site saturation mutagenesis (SSM) library of LuxSit-i. The library was transformed into yeast cells that then underwent protease treatment and sorted to select/enrich for stable variants (that survive the protease treatment) with intact “displayed” protein variants. Individual colonies were then screened on agar plates to identify the brightest clones after addition of the substrate. The top candidates advanced to subsequent rounds of optimization (Supplementary Figure 1). Mutations enhancing brightness were identified at F9, L28, G32, L56, K61, Q64, Q82, A85, H87, W100, R103, and T108 (Supplementary Figure 2).

All the candidate enhancing mutations were combinatorially encoded into a new library, and the best variant from this screen, LuxSit-301 (F9N, Q64L, A85F, H87R, H99L, W100F, T108D, and H113Y, Supplementary Figure 3) exhibited a ∼15-fold improvement in brightness and a reduced proportion of dimeric species (Supplementary Figure 3C).

In the next stage, we screened 56 novel substrates to identify alternatives to DTZ with better solubility, brightness and enhanced glow-time. Luxterazine (LTZ) was identified as the top candidate.

In the next iteration, we sought to optimize LuxSit-301 in the presence of LTZ by using error-prone PCR (EP-PCR) while also aimed to minimize the number of histidine residues in the protein (the parent LuxSit-i, has the remarkably large number of ten total histidines, nine surface-exposed and one in the catalytic pocket). To identify suitable substitutions for the remaining histidine residues, we analyzed the enrichment of mutations after protease treatment of the initial SSM libraries, with enrichment scores reflecting the positive effect of each substitution on protein stability (Supplementary Figure 4). The most favorable (non-Histidine) substitutions were incorporated into a new combinatorial library, together with beneficial mutations identified from the EP-PCR screening. Surprisingly, the top-performing variant from this combinatorial library, LuxSit-2466, was fully devoid of Histidine residues, included the substitution of the catalytic pocket histidine H98Q, a residue that was previously described to be essential for LuxSit function. Our finding establishes that H98Q is functionally advantageous in the context of LuxSit-2466, and remarkably demonstrates that it is possible to generate a highly active version of LuxSit after the complete removal of all histidine residues in the enzyme. LuxSit-2466 exhibited an ∼80-fold increase in brightness over LuxSit-i but also a greater tendency to form oligomeric species (Supplementary Figure 3C). Through three additional rounds of directed evolution, we generated LuxSit-4039, a fully monomeric and highly active enzyme (Supplementary Figure 3D). These final rounds of optimization introduced 14 additional mutations (D23I, V41T, R46V, R55S, L56Q, K68S, Q82R, S92H, T97L, R101K, R103V, D108V, E109A, I114V), unexpectedly one Histidine S91H returned to the original LuxSit sequence for that position; we further confirmed using a single point mutation that the H91 residue contributes favorably to the stability of the enzyme.

In the final step of optimization, using computational protein design three lysine residues were replaced with other residues to generate a lysine-free protein without compromising brightness or stability. This has long been an unresolved but highly sought-after challenge for natural and engineered luciferases, which lose activity in the process of eliminating their surface lysines [9]. The final version of optimized LuxSit was named “LuxSit Pro”, it contains no cysteine or lysine residues, which in turns facilitates its recombinant expression and manufacturing, and furthermore it effectively makes LuxSit Pro an ideal reporter for live-cell applications that rely on independence from the effect of common post-translational modifications (PTM), that often affect the Lysines, such as ubiquitination. Altogether, LuxSit Pro has 27 mutations over LuxSit-i, meaning that ∼23% of the original sequence changed, illustrating the large amount of optimization that was required to deliver the degree of optimization that LuxSit Pro needed to achieve its top performance (Fig. 1A, The sequence of LuxSit Pro is available in Supplementary Table 1).

**Figure 1.**
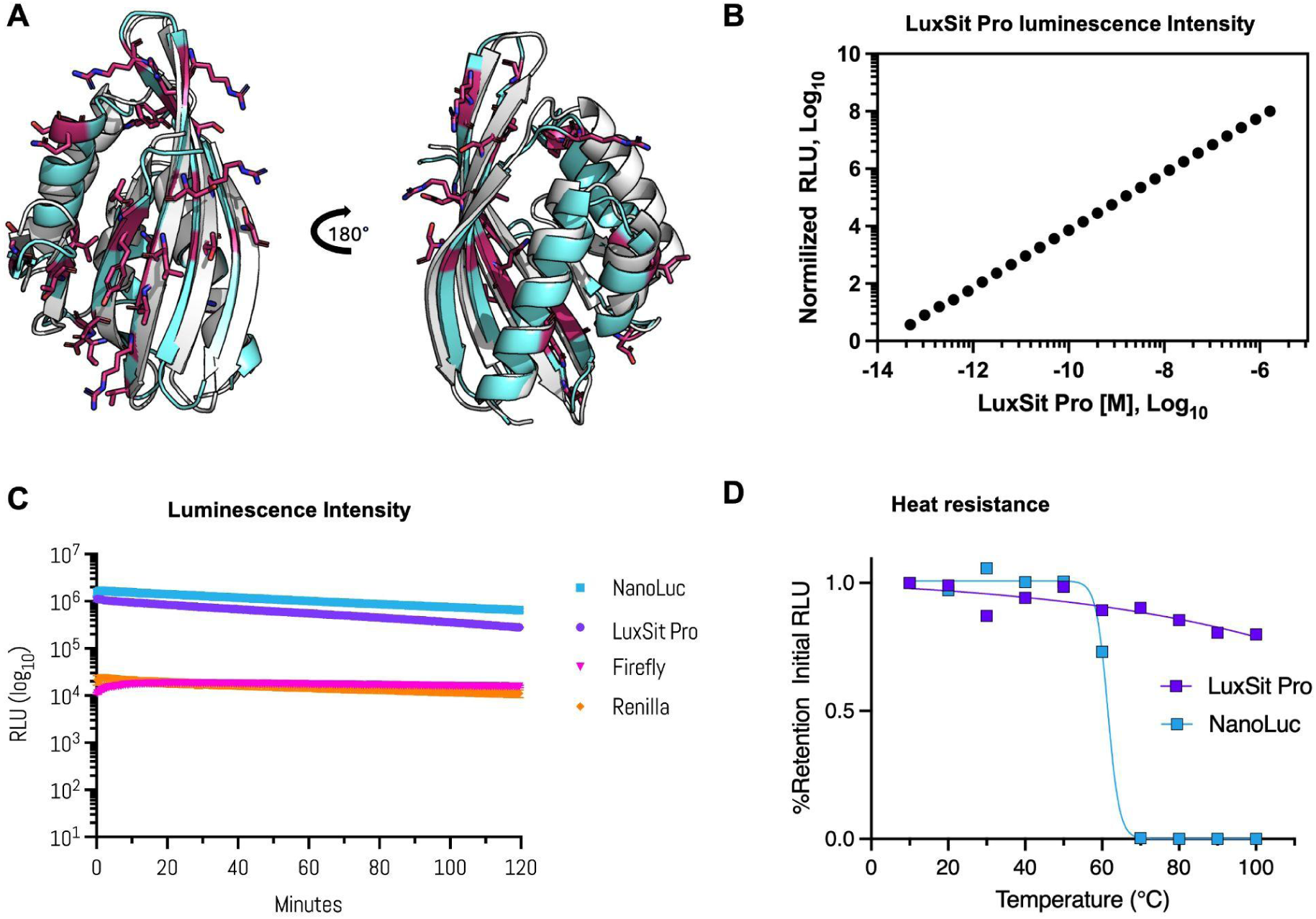
Functional characterization of LuxSit Pro. (A) Structural comparison of LuxSit Pro (cyan) and LuxSit-i (white) AlphaFold2 models, with the 22 mutations highlighted in magenta. (B) Concentration-dependent luminescence of LuxSit Pro. The enzyme exhibits a linear relationship between luminescence intensity and concentration across the tested range, with detectable signal down to 50 femtomolar (fM). (C) Luminescence kinetics in HEK293T cell lysates expressing LuxSit Pro, NanoLuc (furimazine), Firefly luciferase (D-luciferin), or Renilla luciferase (coelenterazine). Measurements were initiated 24 hours post-transfection using manufacturer-recommended substrate conditions. (D) Thermal stability profiles of LuxSit Pro variants and NanoLuc. Protein samples (500 pM) were incubated at 10°C intervals from 10°C to 100°C. Luminescence was recorded 5 minutes after Luxterazine (the LuxSit Pro Substrate) addition using 1c (LuxSit Pro) or furimazine (NanoLuc).

### Performance of the LuxSit Pro system

Monod Bio’s LuxSit Pro is optimized for use with its proprietary substrate Luxterazine and proprietary Assay Buffer (Monod Bio, Cat. #LS0101) [10], producing, in turn, an intense and stable glow-type linear-luminescence-response across several orders of magnitude of the enzyme’s concentration (Fig. 1B). It is ∼150 times brighter than the original LuxSit-i (i.e., comparable in brightness to NanoLuc), and significantly outperforms the brightness of any other known natural luciferase (including Renilla and Firefly luciferases, Fig. 1C). At just 117aa, LuxSit Pro is one of the smallest existing luciferases (13.5 kDa), it recombinantly expresses efficiently in both prokaryotic and eukaryotic systems (Supplementary Figure 5), making it well suited for most in vitro applications, including those that require protein purification (i.e., biosensor manufacturing). LuxSit Pro also demonstrates exceptional thermal stability, it remains fully folded at temperatures up to 95°C (Supplementary Figure 6), and has no dependency on disulfide bridges (since it contains no cysteine residues). Accordingly, LuxSit Pro retains full enzymatic activity after heating to 95°C and cooling back to room temperature (∼20-25°C), whereas under the same experimental conditions, NanoLuc loses function when heated above 60°C (Fig. 1D).

The LuxSit Pro luciferase exhibits a bright peak emission at 490nm (slightly red-shifted compared to LuxSit-i) when used with LTZ (the LuxSit Pro Substrate) (Supplementary Figure 7), and it shows minimal activity with CTZ and commonly used analogs such as FRZ and h-CTZ.

One of the key applications of luciferases is as a reporter in cell experiments. We evaluated the performance of LuxSit Pro across a range of experimental conditions after transfection in HEK293 cells. Transfection efficiency remained high across varying cell densities and DNA input levels (Supplementary Figure 8A-B). In lysing assays LuxSit Pro exhibited excellent tolerance to various cell culture media (Supplementary Figure 8D), fetal bovine serum concentrations (Supplementary Figure 8E), and small amounts of organic solvents (Supplementary Figure 8F). Furthermore, LuxSit Pro retained strong signal intensity across multiple cell-lysate assay formats and sample volumes (from full-well 96-well to 384-well plates, Supplementary Figure 8C), confirming its suitability for high-throughput screening and diverse assay designs. We also confirmed that the assay can be performed without lysing the cells (i.e., in live cell experiments), which indicates that Luxterazine is cell-permeable (results not shown). Future experiments shall address the efficiency and potential improvements to live cell experiments with LuxSit Pro and sPro. Collectively, these results highlight LuxSit Pro as a stable and reliable luciferase for a broad range of experimental applications.

### LuxSit sPro, a reversible two-component complementation system

Building on LuxSit Pro, we developed a two-component complementation system, LuxSit sPro, designed for applications such as bimolecular interaction analysis and biosensing. We tested several options to create a two component system, and determined that splitting LuxSit Pro between positions G104 and N105 (which correspond to the beginning of the C-terminal beta sheet) is an optimum combination for the efficient reconstitution (by proximity) of the enzymatic luciferase activity. The resulting parts are the “large” 103-aminoacid peptide, named aLux (11.8 kDa), and the “small” 12-amino-acid (1.5 kDa) peptide, named bLux (Figure 2A).

**Figure 2.**
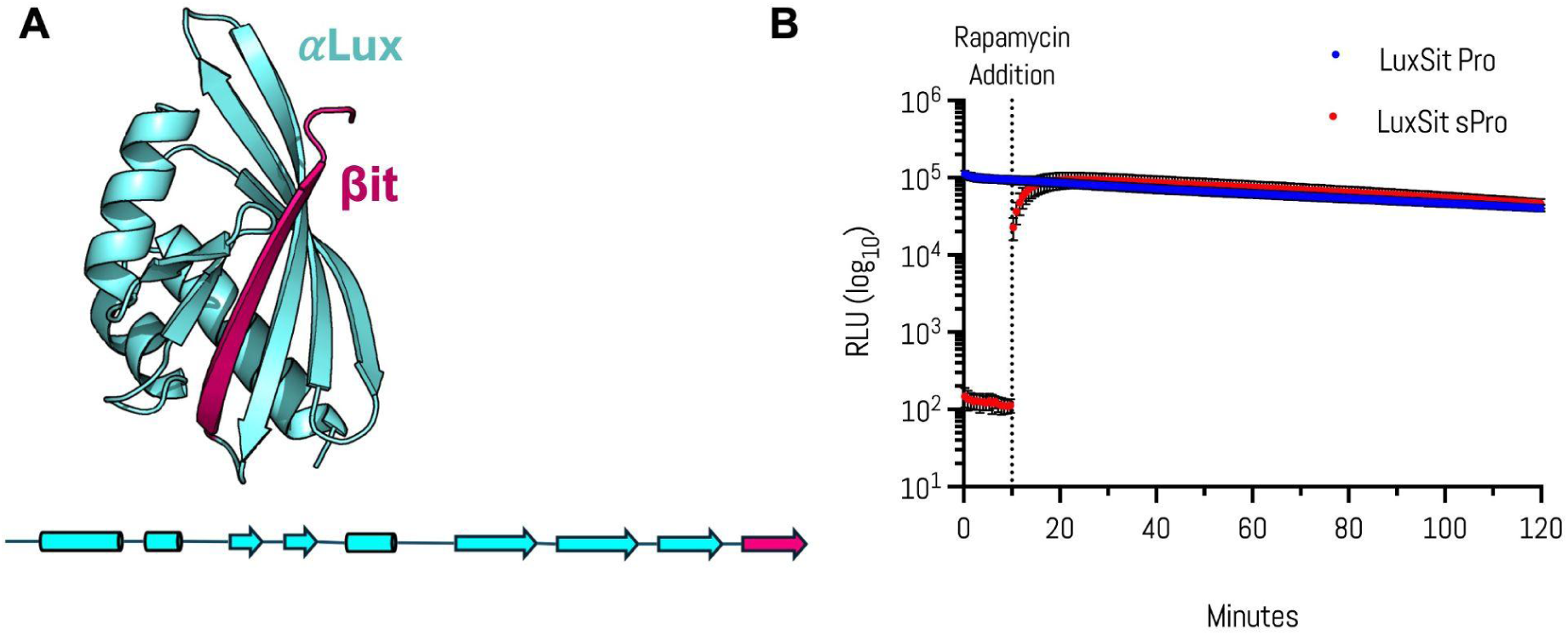
**The LuxSit sPro two-component complementation system.** (**A**) Cartoon representation of the LuxSit sPro structure, highlighting the αLux domain (blue) and βit strand (pink), alongside a schematic 2D model showing the predicted secondary structure organization of the split components. (**B**) To evaluate LuxSit sPro as a molecular interaction reporter, αLux and βit were fused to the rapamycin-binding proteins FKBP and FRB, respectively. Kinetic luminescence assays in mammalian cells compare full-length LuxSit Pro (blue) with the split LuxSit sPro system (red). Baseline luminescence was recorded for 10 minutes prior to the addition of a saturating concentration (100 nM) of rapamycin, followed by continuous monitoring over 120 minutes.

We proceeded to optimize each of the parts for two key properties: stability (to enable seamless plug-and-play genetic fusions without affecting the stability of the fusion partner), and low affinity between components, which is desirable to avoid spontaneous complementation that would interfere with the low-background required for high-sensitivity quantitative applications, such as protein-protein interaction (PPI) quantification.

The initial version of aLux showed a high tendency to aggregate when expressed and purified alone. Therefore, this fragment was optimized through both directed evolution and computational redesign. The best functional variant of aLux was obtained by combining mutations derived from both ProteinMPNN [11] and EP-PCR (note: although we tried, none of the combinatorial libraries from the independent approaches delivered better results). The new version of aLux, named αLux, showed a 10-fold improvement in protein expression, high thermal stability (Tm > 95°C), and it was found to be fully monomeric, making it an ideal “tag” for genetic fusions to proteins of interest (Supplementary Figure 9). To measure the affinity between the two parts of LuxSit sPro, the luminescent activity of αLux was assessed with increasing concentrations of bLux peptide (Supplementary Figure 10). The K_D_ for the reconstitution of aLux and bLux varied from ∼1 to 100µM depending on the bLux sequence/version employed (Supplementary Figure 10). This broad range of affinities confers versatility to the system for different applications. But, for PPI applications we found that the lowest-affinity variant of bLux was the best suited, which we refer from now on as βit. Direct affinity measurements between αLux and βit have a K_D_ of 90µM (±12µM). The final sequences are available on Supplementary Table 1. The LuxSit sPro system demonstrates a bright emission peak at 490 nm (Supplementary Figure 11A), and maintains high specificity for LTZ (Supplementary Figure 11B), and minimal cross-reactivity to CTZ (and its common analogs, such as FRZ and h-CTZ).

We assessed the in vitro performance of αLux and βit using the Rapamycin binding protein pair system (FKBP and FRB). The LuxSit sPro fragments can be fused to either N-terminus or C-terminus of the protein of interest and testing multiple orientations is recommended to achieve optimal activity. We made genetic fusions of the LuxSit sPro fragments to the C-terminus of FKBP and FRB (Supplementary Table 1). At any tested rapamycin concentration, the FKBP-αLux and FRB-βit fusions rapidly reached peak signal (∼5 minutes) upon rapamycin addition, maintaining a sustained response for >120 minutes and achieving up to a 1000-fold increase in luminescence (compared to background). At saturation the brightness of LuxSit sPro is comparable to the full-length LuxSit Pro (Supplementary Figure 12). Dose–response analysis of the rapamycin system resulted in an EC₅₀ of ∼14 nM, which is consistent with previous reports for the rapamycin system (Supplementary Figure 12). Furthermore, the system showed consistent performance/results across a range of cell densities and DNA input levels (Supplementary Figure 13), a highlight of LuxSit sPro robustness across diverse transfection settings. LuxSit sPro efficiently reconstitutes full luciferase activity by proximity, overcoming a common limitation of other split luciferase systems, and it is a reliable and tunable plug-and-play system for studying protein-protein interactions (PPI) on in vitro and cell-based assays (Monod Bio, Cat# SP0101).

To date, we have tested LuxSit Pro in a diverse array of complementation systems in homogeneous assays, such as for the detection of C-reactive protein and hIL-6 (data not-shown), achieving sensitive quantitative detection comparable to state-of-the-art methods.

## Discussion

The computational design of the first de novo luciferase, LuxSit-i [7], opened new avenues for engineering molecular reporters that overcome limitations of natural systems. Since then, Yi *et. al.* recently redesigned LuxSit-i, resulting in NeoLux, which demonstrated a 15-fold increase in luminescence achieved solely through computational methods [8]. Here, we employed a combination of computational protein design, directed evolution and yeast display to improve LuxSit-i activity into a new world-class bio-reporter system. After eight iterative rounds of optimization, we deliver LuxSit Pro and its substrate Luxterazine, which achieves a remarkable ∼150-fold increase in brightness compared to LuxSit-i, highlighting the synergy of combining computational design, directed evolution and substrate redesign to achieve a highly optimized function.

Our fully optimized enzyme, LuxSit Pro, contains 27 mutations compared to LuxSit-i (23% of the sequence). Three mutations – L56Q, H98Q, and W100F – are located within the catalytic pocket. LuxSit Pro catalyzes the oxidation of Luxterazine (LTZ), a novel substrate with improved solubility compared to DTZ, ultimately producing glow-type luminescence. Its brightness enables sensitive detection in a common luminometer plate reader (concentrations as low as 100 fM in a 100µl volume are easily achievable, Fig. 1B). Importantly, LuxSit Pro exhibits orthogonal substrate specificity to NanoLuc (and other luciferase, such Firefly), enabling researchers with a new option for developing multiplexed luminescence based assays by combining the different luciferase systems available in the market.

LuxSit Pro represents a significant advance in luciferase design. Being uniquely free of Lysine and Cysteine residues, our simplified luciferase unlocks new possibilities for site-directed conjugation and allows for robust in vitro studies free from complications caused by common post-translational modifications, such as lysine ubiquitination in the field of Targeted Protein Degradation (TPD). Remarkably, LuxSit Pro achieves peak activity and stability despite its reduced amino acid repertoire, an approach that has proven challenging with other luciferases (as demonstrated by the ∼400-fold signal loss observed when analogous Lysine depletion modifications have been attempted in NanoLuc [9]).

LuxSit sPro, a reversible two-component system built upon LuxSit Pro, enables highly sensitive detection of bi-molecular interactions. Its robust expression, thermal stability, and low intrinsic affinity (90 μM) of the parts allows for rapid and significant reconstitution of luciferase activity by proximity – demonstrated by a ∼1000-fold luminescence increase upon rapamycin addition in a model system.

LuxSit Pro and sPro represent a significant leap forward in luminescent reporter technology, matching and surpassing existing systems in many aspects, that range from its brightness to its hyperstability. The high performance of LuxSit Pro and sPro suggests potential for in vivo applications, possibly by combining Luxterazine (the LTZ), or a variant, with fluorescent molecules to achieve red-shifted emission (i.e., BRET).

As the first de novo-designed protein to enter the commercial market in 2024 (www.monod.bio), and to our knowledge the first de novo-designed enzyme to do so, LuxSit Pro validates the computational protein design paradigm and establishes a new benchmark for protein engineering. LuxSit Pro and LuxSit sPro represent a turning point, demonstrating the power of AI and computational design to generate superior protein tools. While it took a decade for a large corporation to develop NanoLuc, Monod Bio —a small USA startup— delivered LuxSit Pro and sPro to the end-user commercial market in less than two years. We envision a future where computationally engineered proteins routinely surpass those developed through traditional biotechnology, and that future is now within reach.

## Disclaimer

The LuxSit™ assets described in this manuscript are available to the End User Market (B2C and B2B) under a Limited Use Label License (LULL) that requires the sourcing of substrates, assay buffers and/or any other system components directly from Monod Bio, Inc., or its authorized distributors. For more information please visit: https://www.monod.bio.

## Competing interests

RV, JW, JLH, PK, NL, ESF, AQ-R, D-AS, and GJ.L.B. are inventors on a patent application related to the discoveries described in this article. RV, JW, JLH, PK, NL, ESF, AQ-R, and D-AS are employees of Monod Bio, Inc. GJ.L.B. is an advisor to Monod Bio, Inc.

## Materials and methods

### Error-Prone PCR Assembly

Error-prone PCR was performed on LuxSit and its variants using the Agilent Mutazyme II PCR kit (Agilent Technologies). The EP-PCR templates were amplified from vectors containing the respective luciferase genes using KAPA HiFi DNA polymerase (Roche). Reactions were prepared according to the Agilent kit protocol, with template concentrations of 1 and 10, and 25 to 30 cycles to target a mutation frequency of 0–9 mutations per kb.PCR products were purified via gel extraction (QIAquick Gel Extraction Kit, QIAGEN), re-amplified with KAPA HiFi polymerase, and ethanol-precipitated to yield sufficient DNA for transformation. A total of 12 µg of purified DNA was combined with 4 µg of vector pETCON (pre-digested with NdeI and XhoI restriction enzymes) and transformed into electrocompetent EBY100 yeast cells as previously described [12]. The following primers were used for error-prone PCR, initial template amplification, and product re-amplification:

### Forward primer

5′-CGACGATTGAAGGTAGATACCCATACGACGTTCCAGACTACGCTCTGCAG-3′

### Reverse primer

5′-TCCGAACAAAAGCTTATTTCTGAAGAGGACTTGTAATAGAGATCTGATAACAACAGT-3′

### Site Saturation Mutagenesis

Site saturation mutagenesis libraries were generated using NNK OligoPools (Integrated DNA Technologies, IDT). Due to the length of the LuxSit gene (354 bp + vector adapters), which exceeds the 350-bp limit per oligo, the library was split into two halves (177 bp each). Fifty-nine oligos were designed per half, each containing a single NNK degenerate codon at a distinct position.

For the first half oligos, a 50 bp overlap (5′-GAGGAGGCTCTGGTGGAGGCGGTAGCGGAGGCGGAGGGTCGGCTTCGCAT-3′) with the 3′ end of NdeI-XhoI-linearized pETCON3 was added to the 5′ end of the oligo sequences, along with a 20 bp sequence (5′-TCTAAGGACGCGCAACGTGA-3′, positions 178–198) at the 3′ end.

Likewise, for the second half oligos, a 50 bp overlap (5′-GGGGCCTCGAGGGTGGAGGTTCCGAACAAAAGCTTATTTCTGAAGAGGA-3′)wit h the 5′ end of the linearized vector was added to the 3′ end of the oligos, plus a 20 bp sequence (5′-TTGAACGTCTGTTCAGCACC-3′, positions 157–177) at the 5′ end.

Oligos for each half were ordered as separate OligoPools, and unmodified halves with vector overlaps were synthesized as gBlocks (IDT). The OligoPools were re-amplified using KAPA HiFi DNA polymerase and the following primers:

### First half

Forward: 5′-GAGGAGGCTCTGGTGGAG-3′ Reverse: 5′-TCACGTTGCGCGTCCTTAGA-3′

### Second half

Forward: 5′-TTGAACGTCTGTTCAGCACC-3′

Reverse: 5′-TCCTCTTCAGAAATAAGCTTTTGTTCGG-3′

For full-length assembly, 10 ng of the first half OligoPool was combined with 10 ng of the second half gBlock. PCR amplification exploited the 178–198 bp overlap to generate full-length sequences containing vector overlaps and single mutations in the first half. The same process was repeated for the second half OligoPool and first half gBlock (overlap: 157–177 bp).PCR products were gel-purified (QIAquick Gel Extraction Kit, QIAGEN), re-amplified, and ethanol-precipitated. A total of 4 µg of DNA was combined with 1 µg of NdeI/XhoI-digested pETCON vector and transformed into electrocompetent EBY100 yeast cells.

### Combinatorial Library

Combinatorial libraries were constructed to generate all possible combinations of favorable mutations identified through SSM and error-prone PCR. Degenerate codons encoding primarily the selected mutations were introduced into the target luciferase sequence. The modified sequence—including 50 bp overlaps with the pETCON vector at both 5′ and 3′ ends—was divided into 8, 10, or 12 overlapping fragments (30–60 bp each), with degenerate codons excluded from overlapping regions to prevent complementarity issues. Single-stranded primers (IDT) were designed as interleaved forward/reverse pairs, beginning with a forward primer and ending with a reverse primer. Full-length sequences were assembled via PCR by combining 10 ng of each primer fragment. Products of the expected size (∼500 bp) were gel-extracted using the QIAquick Gel Extraction Kit (QIAGEN), re-amplified with KAPA HiFi polymerase, and ethanol-precipitated to yield sufficient DNA for transformation.For yeast transformation, 12 µg of purified DNA was combined with 4 µg of NdeI/XhoI-digested pETCON vector and electroporated into EBY100 cells.

### Yeast Display Screening

SSM, error-prone PCR, and combinatorial libraries were screened by measuring luminescence emission of transformed yeast cells on SD-CAA agar plates. The screening process began with protease treatment to eliminate destabilized variants: libraries were incubated for 10 min at room temperature with diluted protease solutions prepared from stock solutions of 107 µM trypsin and 40 µM chymotrypsin in TBS. The first LuxSit-i SSM library was screened at three different dilutions of the protease stock solution (1/162, 1/273, and 1/729), while all subsequent libraries were screened with a single 1/243 dilution. After protease digestion, cells were washed with PBS containing 1% BSA (PBSF) and labeled with anti-c-Myc-FITC antibody (ICL Lab) to detect surface-expressed luciferases. Fluorescence-activated cell sorting (Sony SH800) was then used to isolate yeast populations that maintained surface expression after protease treatment Sorted cells were grown in SD-CAA liquid media at 30°C, plated on 150 mm SD-CAA agar plates (500–1000 cells per plate), and induced by spraying SG-CAA media on them. Plates were incubated overnight at room temperature, re-sprayed 2–4 hours before imaging, and sprayed with 100 µM substrate. Luminescence was captured (Azure imager) with exposure times ranging from 30 sec (initial library) to 0.5 sec (final library). Parent-sequence colonies were included as background controls. Colonies exceeding parent-sequence luminescence were picked and cultured in 96-deep-well plates (1 mL SD-CAA/well, 30°C). For confirmation, 400 µL of each culture was transferred to 1 mL SG-CAA for overnight induction. Cells (2 million per sample) were resuspended in 90 µL assay buffer, distributed into white 96-well plates (Costar), and sampled with 500 µM substrate. Kinetic luminescence was recorded for 1 hr (Agilent Biotek Synergy H1). Surface expression was quantified via anti-c-Myc-FITC staining (Guava cytometer, Cytek), and emission intensities were normalized to expression levels. Variants with comparable/higher expression and improved normalized activity were selected for sequencing. Plasmids were extracted (Zymoprep-96 Yeast Plasmid Miniprep Kit, Zymo Research), and luciferase genes were PCR-amplified using primers (forward: 5’ - TGACAACTATATGCGAGCAAATCCCCTCAC - 3’; reverse:

AGTCGATTTTGTTACATCTACACTGTTGTTATCAGAT - 3’) for sequencing (Genewiz, Azenta Life Sciences).

### Protein Expression and Purification

The top-performing variants from each optimization cycle were expressed and purified for characterization. DNA sequences encoding the variants - flanked by BsaI restriction sites at both 5’ and 3’ ends - were synthesized as gBlocks (IDT) and cloned into the expression vector LM-627 (Addgene #191551) via Golden Gate assembly. BL21 chemically competent cells (New England Biolabs) were transformed with 5 µL of the assembly reaction and grown overnight at 37°C in 10 mL LB medium supplemented with 50 µg/mL kanamycin. For protein production, cultures were scaled up to 500 mL TB medium with 50 µg/mL kanamycin and grown at 37°C to an OD600 of 0.6-0.8. Expression was induced with 0.5 mM IPTG followed by overnight incubation at 18°C. Cells were harvested, resuspended in lysis buffer (50 mM Tris, 150 mM NaCl), and lysed using a microfluidizer. The clarified lysate was obtained by centrifugation at ∼6,000 × g for 10 min. His-tagged proteins were purified by Ni-Sepharose affinity chromatography (Cytiva). The lysate was loaded onto 1 mL resin pre-equilibrated with wash buffer (50 mM Tris, 150 mM NaCl, 30 mM imidazole). After washing with 4 mL buffer, proteins were eluted with 3 mL elution buffer (50 mM Tris, 150 mM NaCl, 300 mM imidazole). Final purification was achieved by size-exclusion chromatography using a Superdex 75 column (Cytiva) on an ÄKTA pure system (Cytiva), with fractions eluting at the expected volume collected for analysis. Protein purity was verified by SDS-PAGE analysis, with fractions showing >90% purity pooled for downstream applications.

### Biophysical characterization

Purified proteins were characterized for thermal stability and polydispersity. Thermal melt assays were performed in PBS (0.2 mg/mL protein) using a temperature ramp from 20°C to 95°C at 1°C/min. Circular dichroism (CD) signals were monitored at 222 nm in 0.1 cm pathlength quartz cuvettes with a MOS-500 spectropolarimeter (Biologic). Raw data (millidegrees) were converted to mean residue ellipticity (MRE) and plotted as a function of temperature. Thermal recovery was assessed using an UNCLE system (Unchained Labs) with samples at 0.2 mg/mL in PBS. The temperature program followed an oscillating profile where the sample was returned to 20°C for 300 s equilibration before each new target temperature. The sequence progressed in 15°C increments (20°C→35°C→20°C→50°C→20°C→65°C→20°C→80°C→20°C→95°C) followeD by 15°C decrements (95°C→20°C→80°C→20°C→65°C→20°C→50°C→20°C→35°C→20°C), completing nine full cycles. Protein refolding competence was monitored via static light scattering (SLS) at 266 nm. Polydispersity analysis was conducted by SEC-MALS using a MiniDAWN multi-angle light scattering detector and Optilab refractive index detector (Wyatt Technology). Protein samples (∼1 mg/mL in PBS) were filtered (0.05 µm) and injected (100 µL) onto a Superdex 75 column (Cytiva) coupled to an HPLC Infinity II 1260 system. Data were processed using Astra software (Wyatt) to determine molecular weights and polydispersity indices.

### Luminescence Signal Characterization of Full-Length Enzymes

Luminescence kinetics were measured following the instructions in the LuxSit Pro manual (Monod Bio, Cat# LS0101). In brief, purified LuxSit variants (555 pM in assay buffer) were dispensed (90 µL/well) into white 96-well plates (Corning). Reactions were initiated by adding 10 µL of 1 mM substrate (final concentrations: 50 µM substrate, 500 pM protein). For initial variants (including LuxSit-i and MBIO-301), DTZ substrate in PBS was used; while all subsequent optimized variants employed Luxterazine (the LuxSit Pro Substrate) in assay buffer. Luminescence was recorded (Agilent Biotek Synergy H1) for 1 hr at 25°C (gain: 100, shaking every 2 min before readings). For quantitative comparison of the emission intensities, we performed kinetic measurements on serially diluted samples of LuxSit Pro, LuxSit-i, and NanoLuc. Proteins were diluted 2-fold from 500 pM to 3.9 pM in their respective buffers. Following Luxterazine addition (50 µM final concentration), luminescence was monitored for 1 hr. The signal intensity at ∼5 min was plotted against protein concentration. Intensity differences were calculated from signals at the 500 pM concentration. Protein stability was assessed by monitoring luminescence retention after thermal challenge. Enzyme solutions (200 pM) were heated in 10°C increments (10-100°C, 5 min per step) in a PCR cycler, then cooled to room temperature before adding 50 µM substrate. The 5-minute post-initiation luminescence values were normalized to untreated controls (set as 100% signal output) to generate thermal resistance profiles.

### LuxSit Pro Split Reconstitution Assays

The LuxSit sPro luminescent assays were performed following the instructions in the LuxSit sPro manual (Monod Bio, Cat # SP0101). To evaluate complementation of the split LuxSitPro system, we measured luminescence recovery upon combining the purified αLux fragment with synthetic βit peptide. While αLux was expressed and purified as described previously, the short βit peptide was synthesized commercially (GenScript) and delivered as a lyophilized powder. The peptide was reconstituted in DMSO to prepare a 4 mM stock solution. For the reconstitution assay, we prepared serial dilutions of βit peptide (1.6 mM to 0 mM in 2-fold steps) in the assay buffer. Each dilution (10 μL) was mixed with 80 μL of 5 nM αLux solution in white 96-well plates, yielding final βit concentrations ranging from 160 μM to 0 μM. Following a 10-minute incubation at room temperature to allow complex formation, reactions were initiated by adding 10 μL of 500 μM substrate (50 μM final concentration). Luminescence kinetics were recorded as previously described, with the signal intensity at 5 minutes post Luxterazine addition used for analysis. The dose-response relationship was analyzed by fitting the data to a one-site binding model using GraphPad Prism software to estimate the αLux-βit binding affinity.

### Rapamycin Binding Detection via LuxSit Pro Reconstitution in Mammalian Cells

Transfection complexes were prepared prior to cell plating to ensure proper formation. For each well, 10 µL of transfection mix containing DNA and transfection reagent (Lipofectamine, ThermoFisher #18324012) was prepared according to manufacturer protocols. The DNA-reagent mixture was vortexed and incubated at room temperature for ≥15 min. HEK293T cells were counted and resuspended at 2×10⁵ cells/mL in fresh medium. Cells (100 µL/well, 2×10⁴ cells/well) were plated in 96-well plates, followed by addition of 10 µL transfection mix per well with gentle swirling to ensure mixing. Transfected cells were maintained at 37°C/5% CO₂ for 24 hr. After incubation, plates were centrifuged (300 × g, 5 min) and media aspirated. Cells were washed with 100 µL DPBS, centrifuged again (300 × g, 5 min), and lysed with 20 µL Passive Lysis Buffer (Promega, #E1941, 10 min incubation). Substrate (120 µM in assay buffer, 100 µL/well) was added and background luminescence recorded for 10 min. Rapamycin response was measured after adding 10 µL/well of compound, with luminescence recorded using standard plate reader settings.

### Reconstitution of LuxSit™ Pro Substrate

Lyophilized LuxSit Pro Substrate (Monod Bio, Inc) and the accompanying Reconstitution Buffer were removed from –20°C storage and allowed to equilibrate to room temperature prior to use. Once equilibrated, the Reconstitution Buffer was gently mixed by inversion. Subsequently, 2.5 mL of Reconstitution Buffer was added directly to the vial containing the lyophilized LuxSit Pro Substrate. The solution was gently inverted until the substrate was fully dissolved, yielding a 5 mM stock solution. To minimize photodegradation, all handling of the substrate was performed with limited exposure to light. The reconstituted substrate was stored at –20°C when not in use. To reduce the number of freeze-thaw cycles, the stock solution was aliquoted into smaller volumes. For reference, preparation of Assay Reagent sufficient for a full 96-well plate requires 240 µL of the 5 mM stock solution.

### Preparation of Assay Reagents

Assay Buffer from the LuxSit Pro ki (Monod Bio, Cat #LS0101) was equilibrated to room temperature prior to use by thawing either at 4°C or at room temperature. Once fully thawed, the buffer was gently mixed by inversion. The reconstituted LuxSit Pro Substrate stock was also equilibrated to room temperature in the dark prior to use. The Assay Reagent was prepared fresh for each experiment by diluting one volume of 5 mM LuxSit Pro Substrate stock with 50 volumes of Assay Buffer (1:50 dilution). For example, to generate 10 mL of Assay Reagent, 240 µL of the 5 mM stock substrate was diluted into 9.76 mL of Assay Buffer. The final Assay Reagent was protected from light and used promptly. Performance testing indicated the Assay Reagent retained up to 75% of its peak luminescence for up to three hours after preparation when compared to freshly prepared reagent.

### Transfection Optimization with Varying Cell Densities

To evaluate the effect of cell density on transfection efficiency using the LuxSit s-Pro PPI system, HEK293T cells were seeded at varying densities in 96-well plates and transfected with 20 ng of each LuxSit s-Pro PPI plasmid per well. Following an overnight incubation at 37 °C with 5% CO₂, cells were lysed in lysis buffer for 15 minutes at room temperature. Subsequently, LuxSit Pro Substrate diluted in the supplied Assay Buffer was added to each well. Luminescence kinetics were recorded over time, and data were plotted to compare signal dynamics across different initial cell densities.

### Transfection Optimization with Varying DNA Amounts

To assess the influence of DNA input on the assay performance, 10,000 HEK293T cells per well were seeded in a 96-well plate and transfected with varying amounts of LuxSit s-Pro PPI plasmid DNA. After overnight incubation, cells were lysed for 15 minutes at room temperature. LuxSit Pro Substrate, diluted in Assay Buffer, was then added to each well. Luminescence was measured, and the maximum signal following Rapamycin addition was quantified. The data represent the average luminescence (n = 4) expressed as fold-change over background.

## Supplementary Figures

**Supplementary Table 1.**
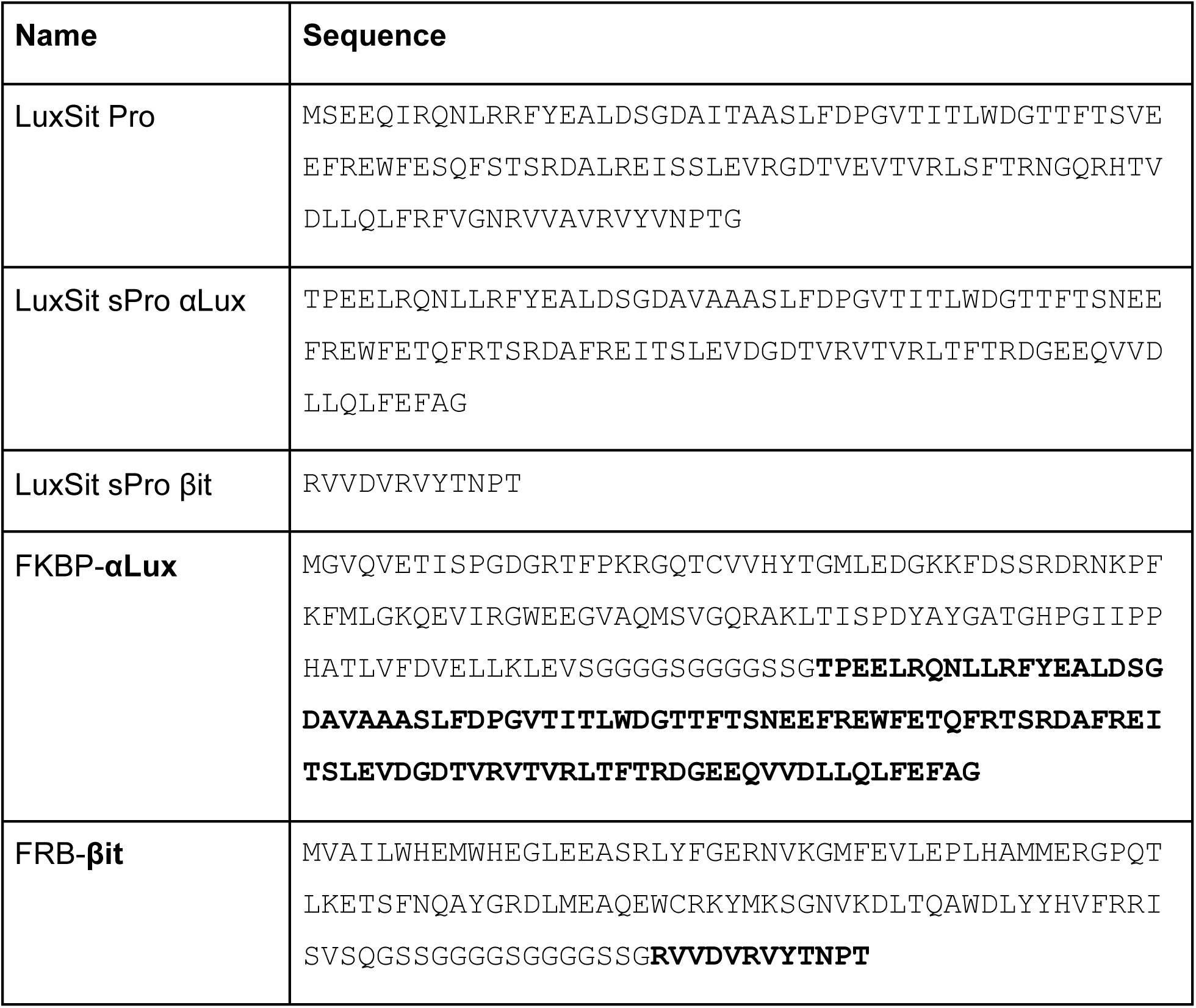
Sequences for LuxSit Pro and the LuxSit sPro systems.

**Supplementary Figure 1.**
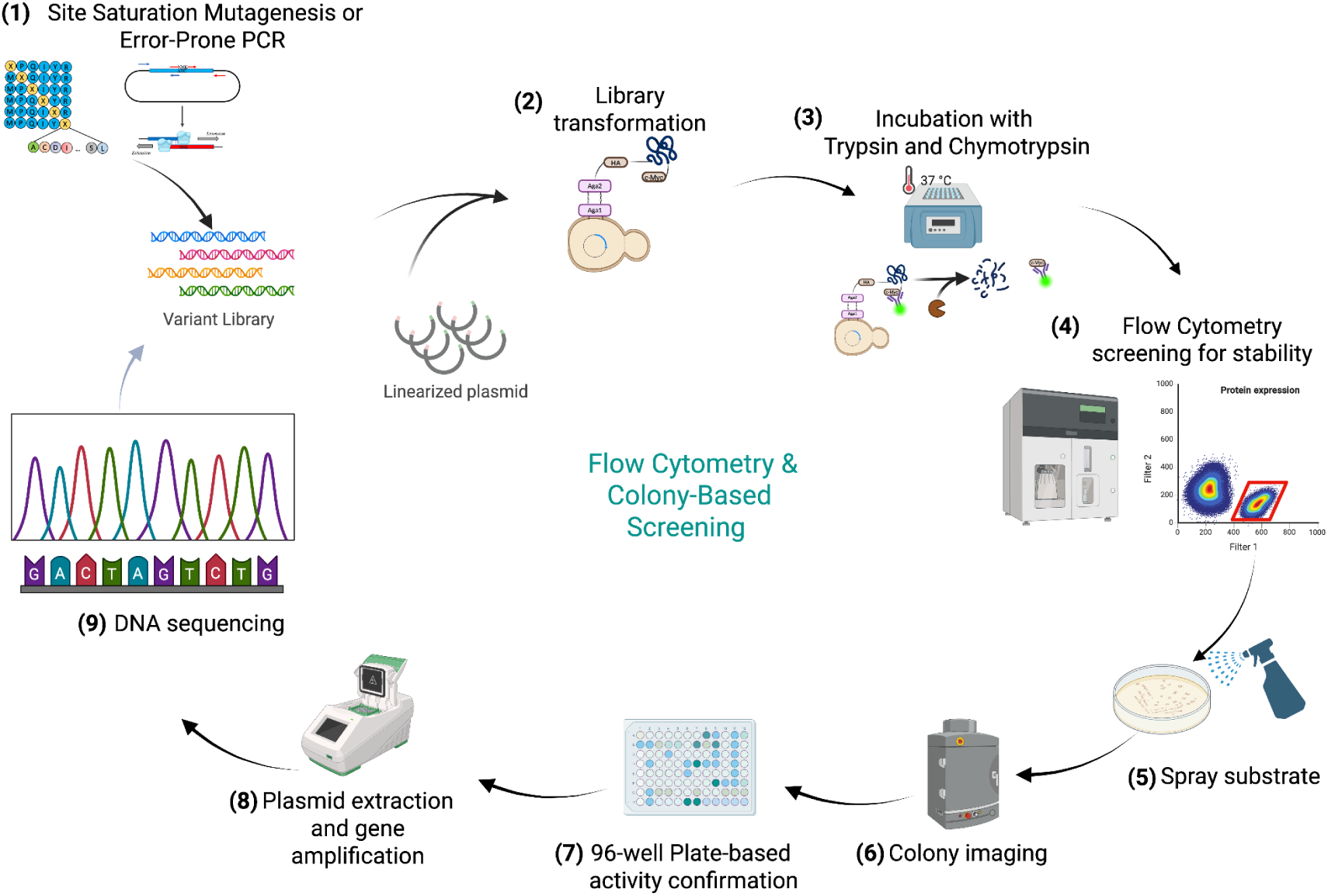
Workflow for iterative optimization of LuxSit variants. Each round of directed evolution followed these sequential steps: (1) Library generation via site-directed mutagenesis or error-prone PCR; (2) Co-transformation of EBY100 yeast with library DNA and linearized pETCON plasmid via electroporation; (3) Protease selection using optimized trypsin/chymotrypsin concentrations to eliminate unstable variants; (4) Fluorescence-activated cell sorting (FACS) to isolate cells expressing protease-resistant variants; (5) Plate-based screening on 150 mm SDCAA agar, with protein expression induced by SGCAA media spraying followed by substrate (100 µM) application for luminescence detection; (6) Imaging of luminescent colonies using a CCD camera system immediately after Luxterazine addition; (7) Validation of top-performing colonies through liquid culture assays measuring expression-normalized luminescence in 96-well plates; (8) Plasmid extraction and gene amplification by PCR from selected variants; (9) Nanopore or Sanger sequencing of improved clones. Confirmed variants served as templates for subsequent optimization rounds, with purified protein validation between cycles.

**Supplementary Figure 2.**
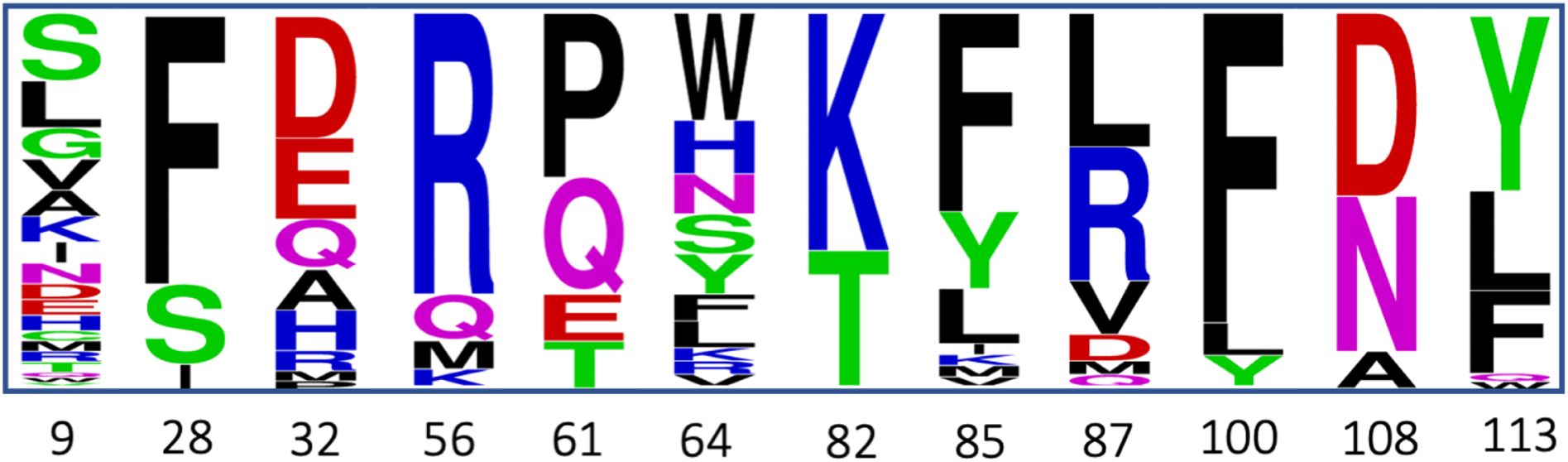
Sequence analysis of first-round optimization variants. The sequence-logo displays amino acid positions and substitutions enriched in improved variants following site-directed mutagenesis and selection. Residue numbering corresponds to the LuxSit-i parent sequence, with letter height proportional to mutation frequency at each position.

**Supplementary Figure 3.**
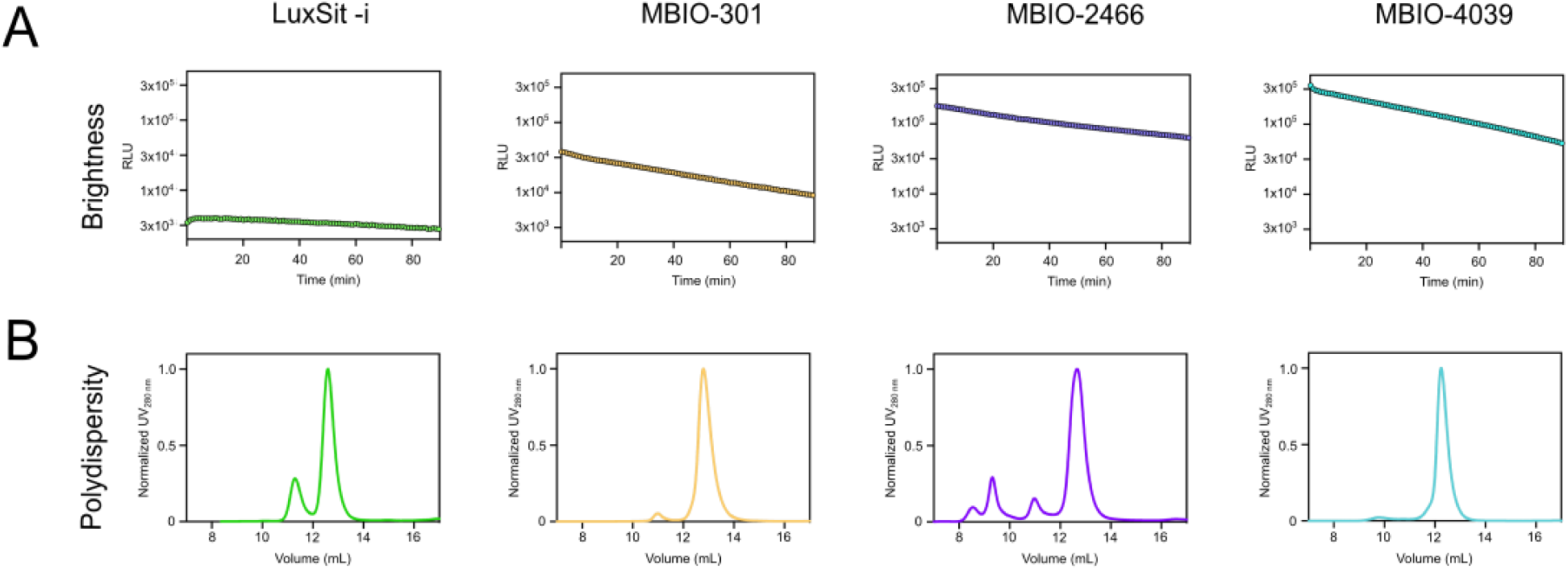
Directed evolution optimization of LuxSit-i. (A) Luminescence kinetics of LuxSit-i (green) versus optimized variants (LuxSit-301: yellow; LuxSit-2466: purple; LuxSit-4039: cyan) measured under identical experimental conditions. (B) SEC-MALS elution profiles (normalized UV 280 nm traces) comparing the oligomeric states of LuxSit-i (green) and optimized variants at ∼1 mg/mL protein concentration. All structural and biochemical characterizations were performed in parallel to enable direct comparison.

**Supplementary Figure 4.**
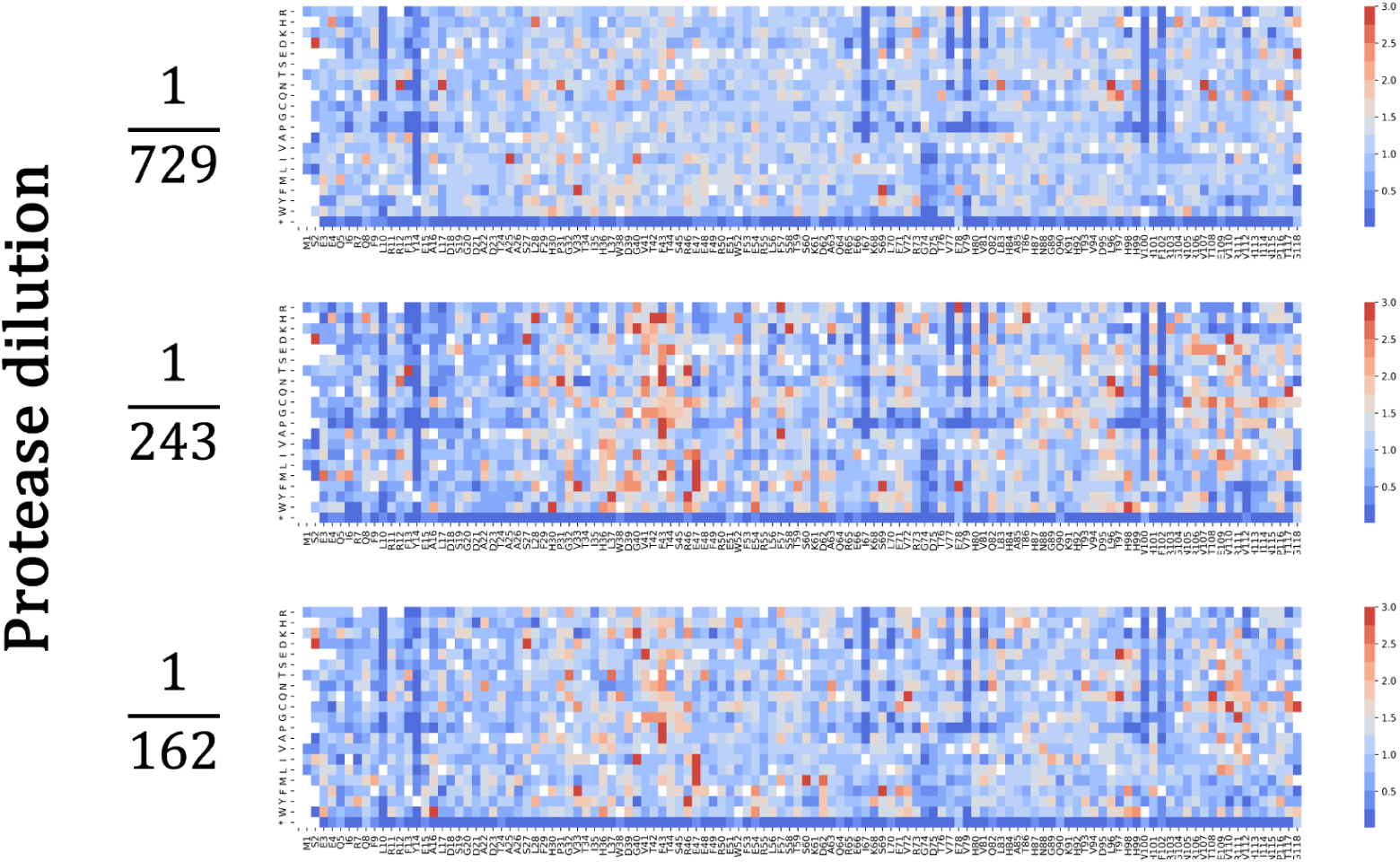
Protease resistance profiling of site-saturation mutagenesis libraries through enrichment analysis. Heat maps visualize mutation enrichment values (Y-axis) across LuxSit amino acid positions (X-axis), with a color gradient representing enrichment levels from lowest (blue) to highest (red). Three protease conditions are shown: 1/729, 1/243, and 1/162 dilutions of stock protease solutions (107 µM trypsin and 40 µM chymotrypsin). The analysis reveals minimal enrichment for mutations at core hydrophobic residues L10, Y14, I67, V79, V100, and F102, indicating strong conservation of these structurally critical positions. At protease concentrations of 1/243 and 1/162 dilutions, two distinct regions show pronounced mutation enrichment corresponding to secondary structure elements: the first spanning β-strand 2 (residues G40-E47) and the second encompassing β-strand 6 (residues R106-G118). These patterns demonstrate both the essential role of the hydrophobic core in maintaining protein stability and the potential for surface-exposed β-strands to accommodate stabilizing mutations under selective pressure.

**Supplementary Figure 5.**
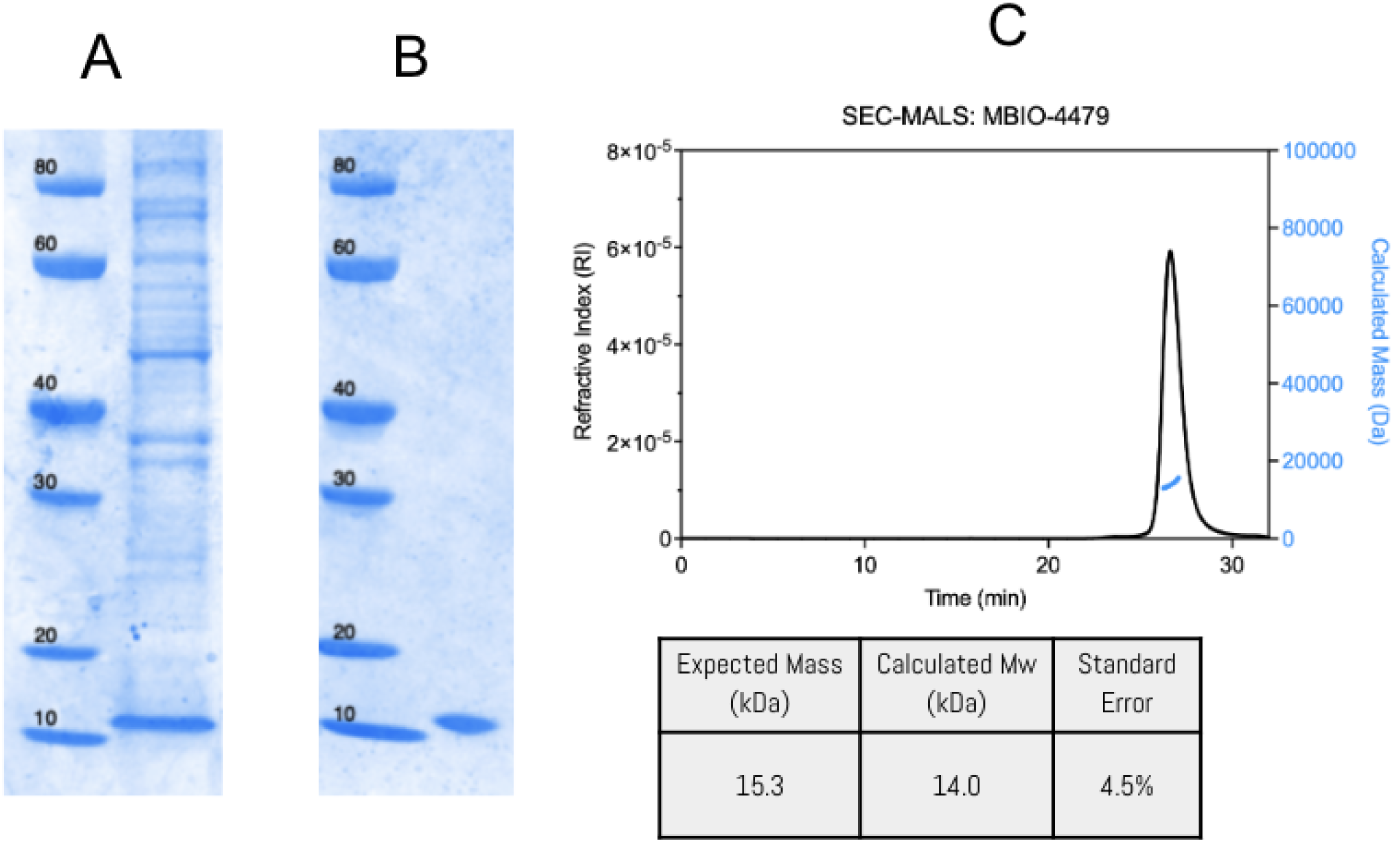
Bacterial expression and purification of LuxSit Pro. (A) SDS-PAGE gel of whole-cell expression, stained with Coomassie blue, showing robust expression of LuxSit Pro. (B) Purified LuxSit Pro following immobilized metal affinity chromatography (IMAC) and size-exclusion chromatography (SEC), yielding 56 mg of protein per liter of bacterial culture. (C) SEC-MALS analysis confirming that LuxSit Pro is >95% monomeric and elutes with the expected molecular mass corresponding to a monomer.

**Supplementary Figure 6.**
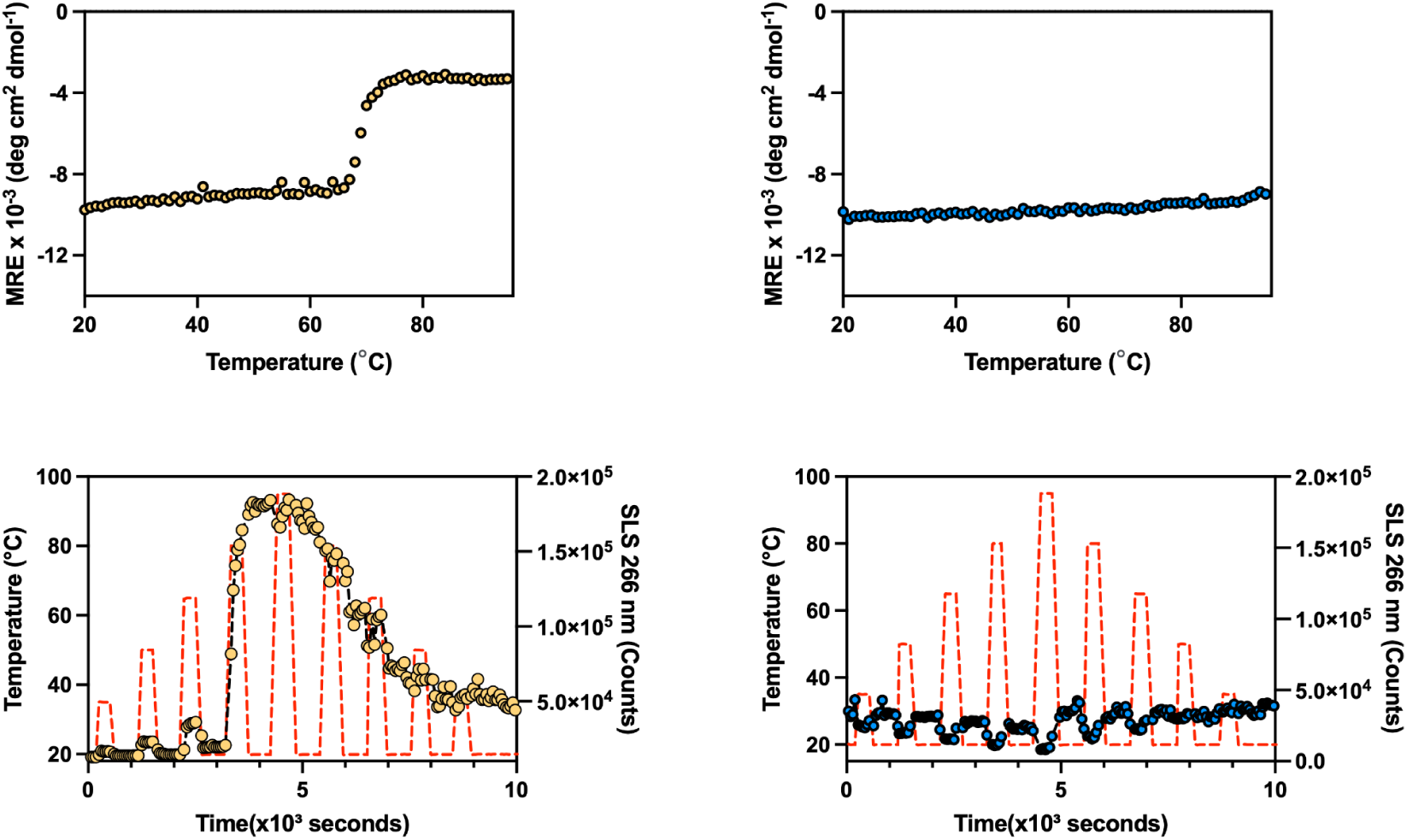
Comparative thermal stability analysis of NanoLuc and LuxSit Pro. Upper panels: Circular dichroism (CD) thermal unfolding curves showing a distinct transition with a melting temperature of 69°C for NanoLuc (left, yellow), while LuxSit Pro (right, blue) maintains its folded state without an observable transition. Lower panels: Thermal recovery assays monitoring static light scattering (SLS) signals during repeated heating-cooling cycles (red dashed lines indicate temperature transitions). NanoLuc (left, yellow) exhibits a substantial increase in SLS signal at ∼80°C, indicative of aggregation, while LuxSit Pro (right, blue) maintains baseline SLS levels throughout all thermal cycles, demonstrating aggregation resistance.

**Supplementary Figure 7.**
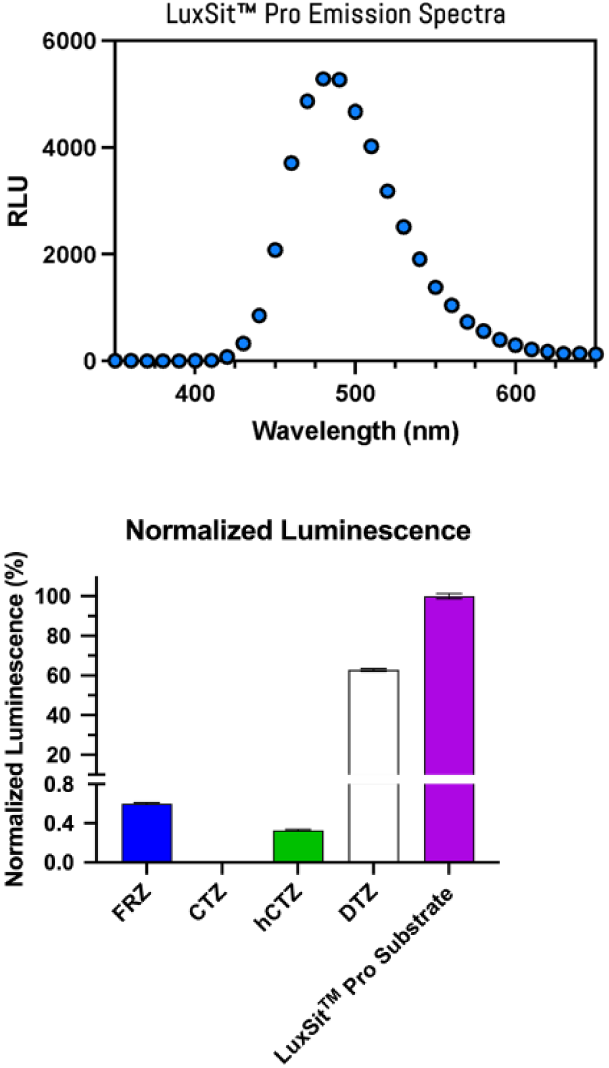
Emission spectra and substrate compatibility of LuxSit Pro luciferase. (A) Emission spectrum of LuxSit Pro luciferase, showing a peak emission wavelength at 490 nm. (B) Substrate compatibility profile of LuxSit Pro luciferase. Various substrates – Luxterazine (the LuxSit Pro Substrate), FRZ, CTZ, hCTZ, and DTZ – were each diluted to 60 µM and mixed 1:1 with purified LuxSit Pro protein at 500 pM. Luminescence was measured immediately after Luxterazine addition. Data represent the average luminescent signal (n = 3), normalized to the signal obtained with Luxterazine (the LuxSit Pro Substrate).

**Supplementary Figure 8.**
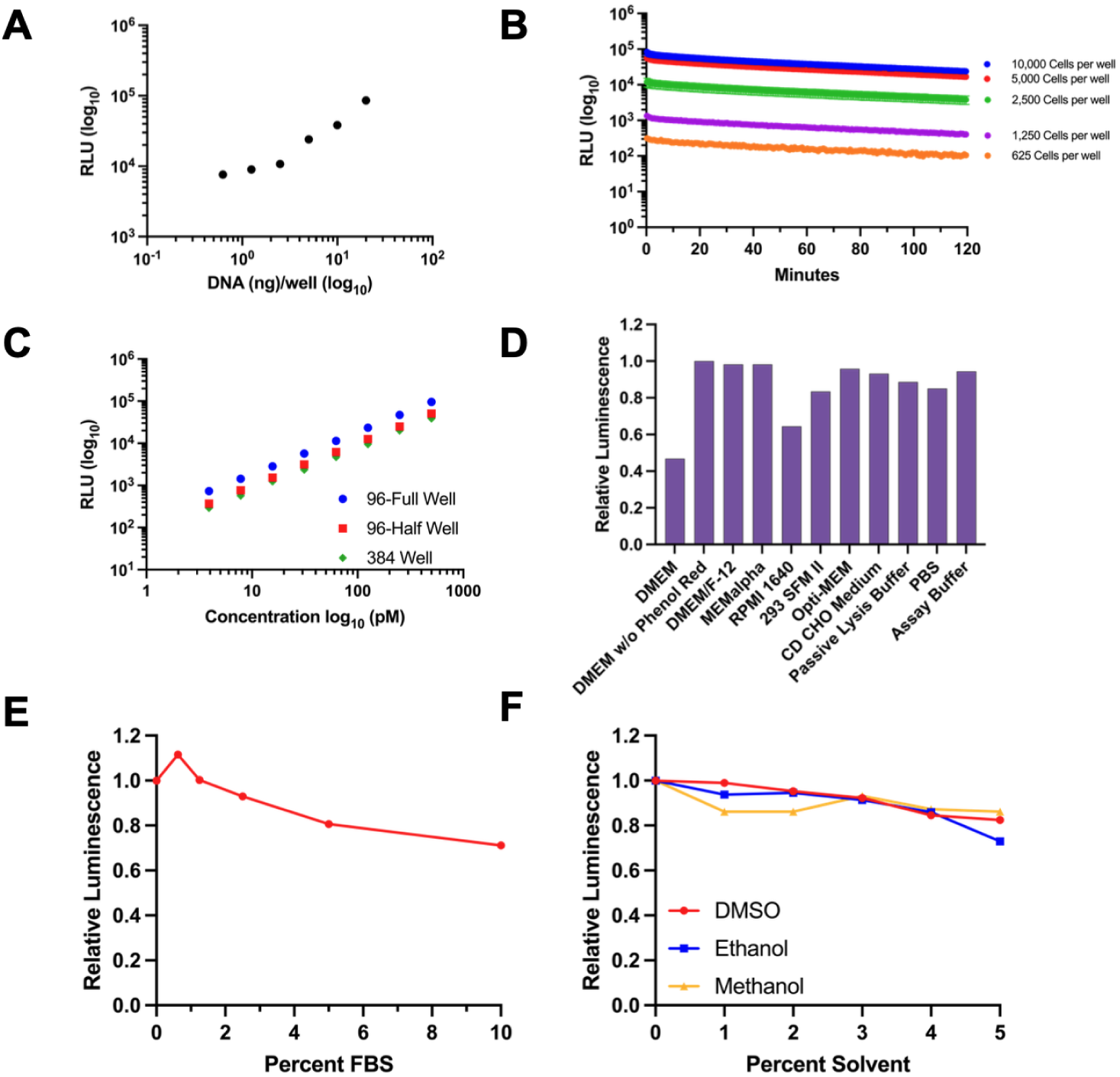
Factors Impacting LuxSit Pro Performance. The impact of various experimental conditions on LuxSit Pro luciferase activity was evaluated using either transfected HEK293T cells (Panels A–B) or purified protein (Panels C–F). (A) Effect of DNA transfection amount on luminescence signal. (B) Influence of initial cell plating density. (C) Comparison of different plate formats. (D) Effect of cell culture media type. (E) Influence of fetal bovine serum (FBS) concentration in the reaction buffer. (F) Tolerance of LuxSit Pro to various solvents at different concentrations.

**Supplementary Figure 9.**
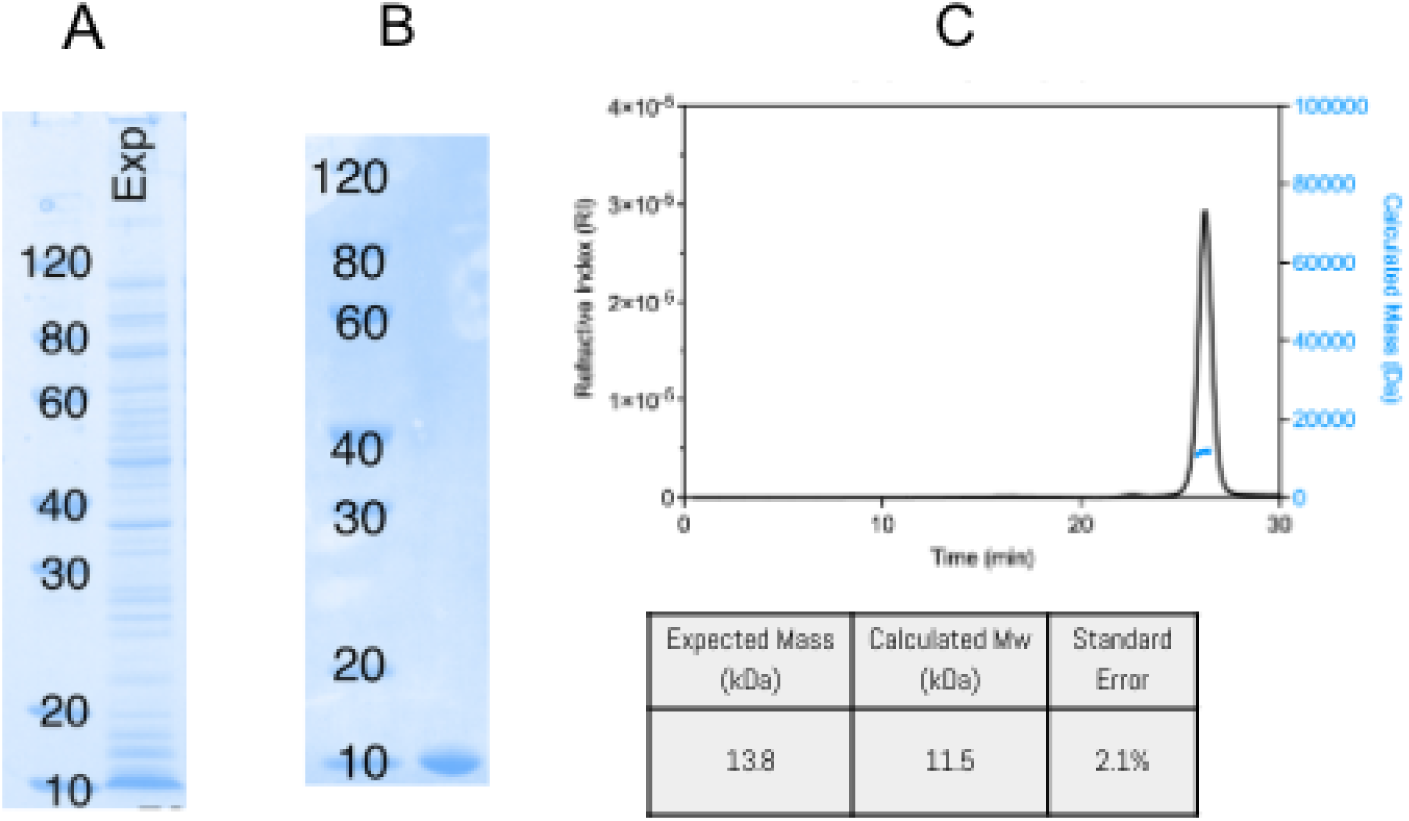
Bacterial expression, purification, and stability of LuxSit sPro αLux. (A) SDS-PAGE gel of whole-cell expression, stained with Coomassie blue, showing robust expression of LuxSit sPro αLux. (B) SDS-PAGE gel of purified LuxSit sPro αLux following immobilized metal affinity chromatography (IMAC) and size-exclusion chromatography (SEC), yielding 43 mg of protein per liter of bacterial culture. (C) SEC-MALS analysis confirming that LuxSit sPro αLux is >95% monomeric and elutes with the expected molecular mass corresponding to a monomer.

**Supplementary Figure 10.**
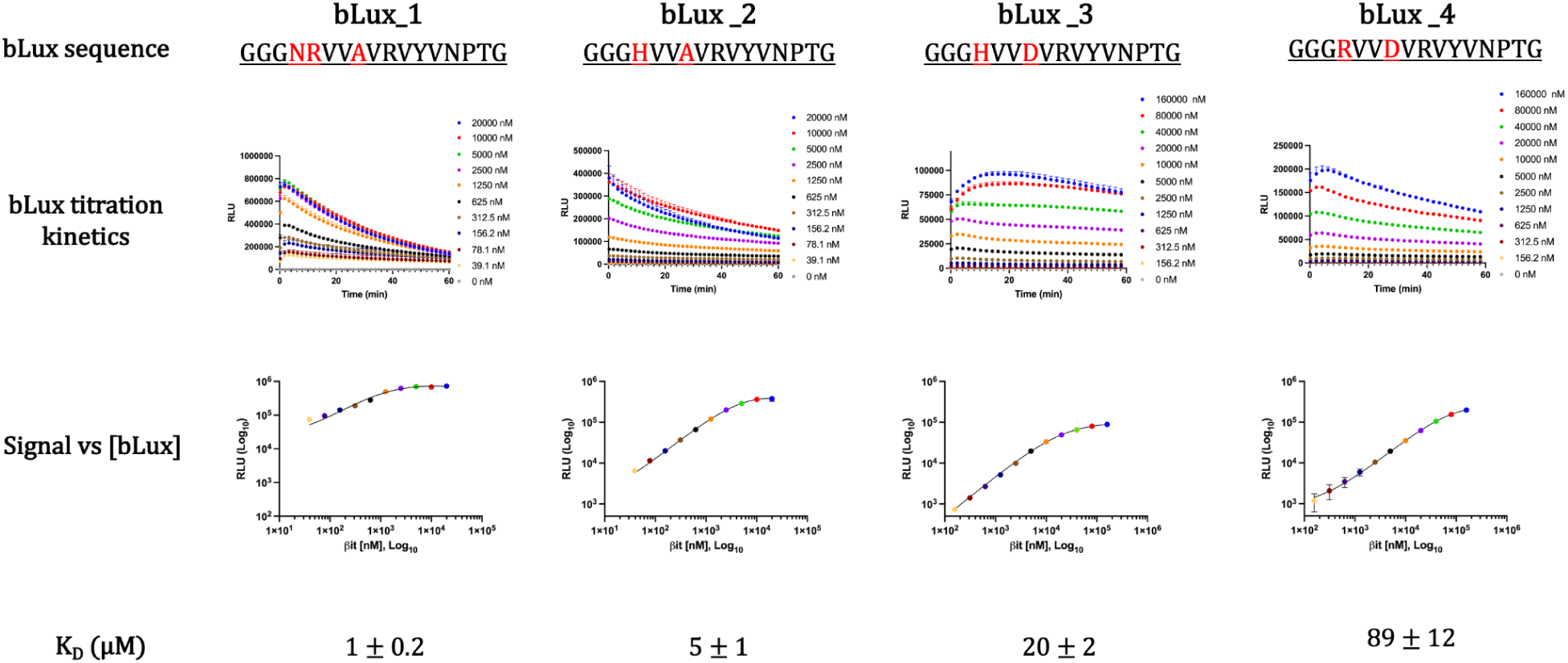
Binding affinity analysis of αLux-bLux interactions. Serial dilutions of different bLux sequences (amino acid differences shown in red) were titrated against αLux (0-20 μM or 0-160 μM range). Luminescence was recorded, and the signal at ∼3 minutes after Luxterazine addition was plotted against bLux concentration. Data were fit to a single-site binding model: Y = Bmax*X/(Kd + X) + NSX + Background, where X is βit concentration, Bmax is maximum specific binding, NS represents nonspecific binding, and Background is baseline signal. Calculated dissociation constants (Kd) are displayed for each bLux variant. The final LuxSit sPro βit is bLux_4.

**Supplementary Figure 11:**
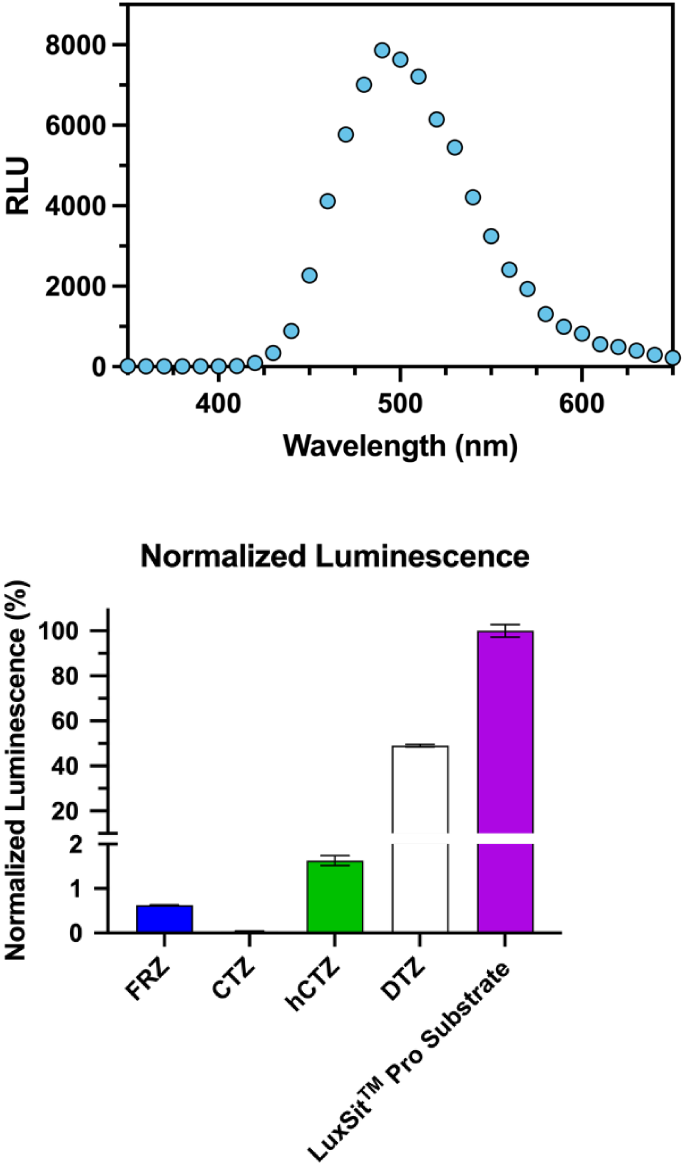
(A) Light emission spectrum of reconstituted LuxSit sPro luciferase with Luxterazine (the LuxSit Pro Substrate), showing a peak emission wavelength at 490 nm. Active enzyme was generated by rapalmycin-induced complementation of FKBP-αLux and FRB-βit fusion proteins, as described in Figure 2, using 100 nM rapamycin and a 120-minute incubation. (B) Substrate compatibility profile of LuxSit sPro luciferase. Luxterazine (LTZ, LuxSit Pro Substrate) and alternative substrates—FRZ (furimazine), CTZ (coelenterazine), hCTZ (h-coelenterazine), and DTZ (diphenylterazine)—were each diluted to 60 µM and mixed 1:1 with purified, reconstituted LuxSit sPro protein at 500 pM. Luminescence was measured immediately after Luxterazine addition. Data represent the average luminescent signal (n = 3), normalized to the signal obtained with Luxterazine (the LuxSit Pro Substrate).

**Supplementary Figure 12.**
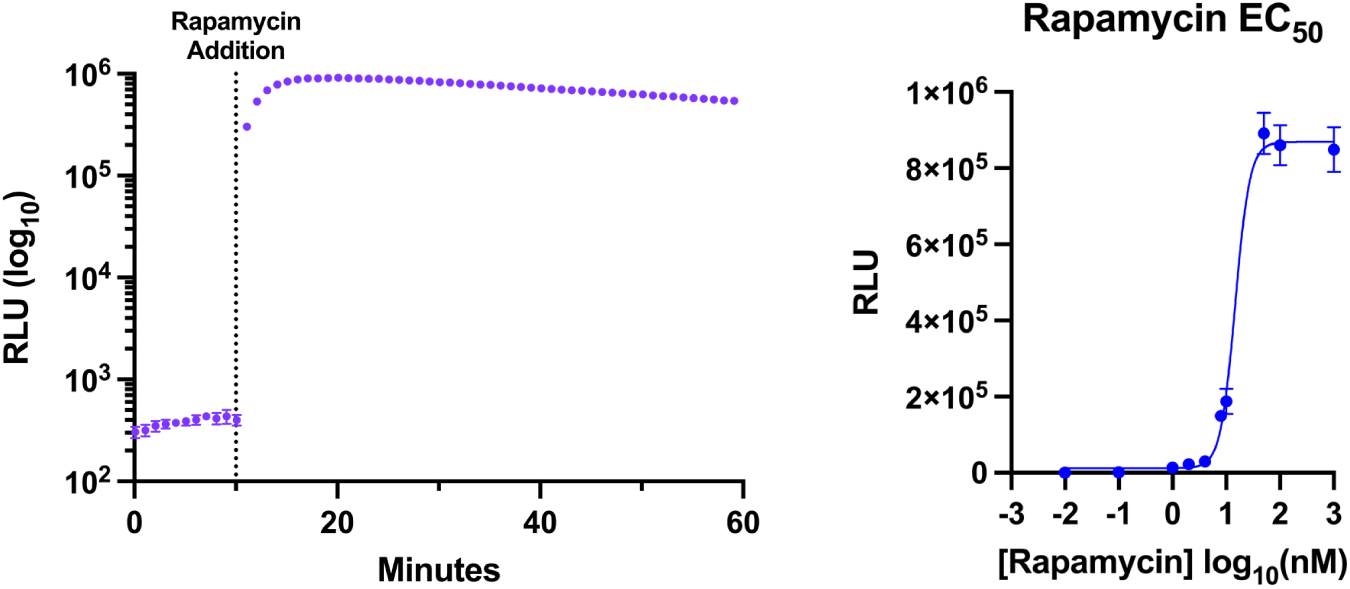
Detection and rapamycin responsiveness of the LuxSit sPro PPI system in lysed mammalian cells. (A) HEK293T cells were transfected with LuxSit sPro plasmids and incubated for 24 hours. Cells were lysed using 1× Passive Lysis Buffer (Promega, Cat. No. E1941), followed by addition of Luxterazine (the LuxSit Pro Substrate) Reagent. Bioluminescence was recorded using a standard plate reader. After a 10-minute baseline read, 100 nM rapamycin was added to induce complementation. Relative luminescence units (RLU) are shown as the mean of triplicate transfections, with error bars representing standard deviation. Peak signal was observed 10 minutes after rapamycin addition. (B) Determination of rapamycin EC₅₀ for LuxSit sPro PPI control plasmids. HEK293T cells were transfected in 96-well plates with LuxSit sPro PPI control DNA and incubated overnight. Cells were lysed for 15 minutes, followed by addition of Luxterazine diluted in LuxSit Pro Assay Buffer. After a 10-minute baseline read, serial dilutions of rapamycin were added. Maximum luminescent signal over 1 hour (n = 3) is plotted for each concentration. The calculated EC₅₀ was 14 nM.

**Supplementary Figure 13.**
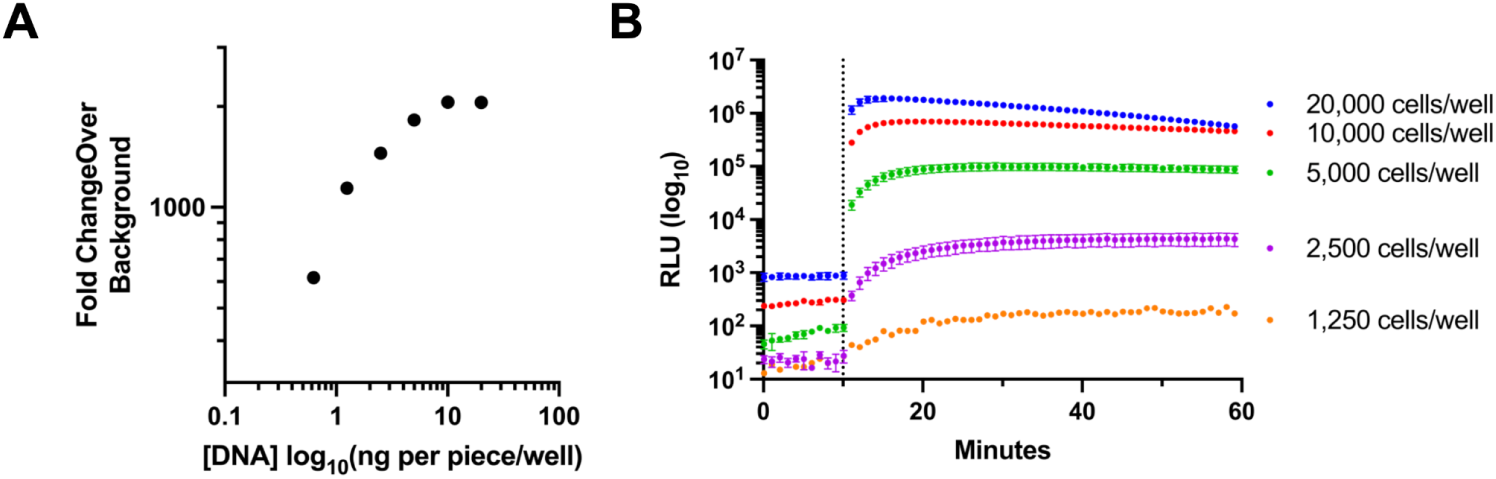
Impact of cell number and DNA transfection amount on LuxSit sPro performance in live-cell experiments. Two conditions were tested to examine their impact on LuxSit sPro luciferase performance using HEK293T transfected cells: (A) amount of DNA transfected, (B) number of plated cells.

## References

1. Delroisse J, Duchatelet L, Flammang P, Mallefet J. Leaving the Dark Side? Insights Into the Evolution of Luciferases. Front Mar Sci. 2021;8: 673620.

2. Zambito G, Chawda C, Mezzanotte L. Emerging tools for bioluminescence imaging. Curr Opin Chem Biol. 2021;63: 86–94.

3. Yeh H-W, Ai H-W. Development and Applications of Bioluminescent and Chemiluminescent Reporters and Biosensors. Annu Rev Anal Chem (Palo Alto Calif). 2019;12: 129–150.

4. Hall MP, Unch J, Binkowski BF, Valley MP, Butler BL, Wood MG, et al. Engineered Luciferase Reporter from a Deep Sea Shrimp Utilizing a Novel Imidazopyrazinone Substrate. 2012 [cited 17 Jul 2025]. doi:10.1021/cb3002478

5. Dixon AS, Schwinn MK, Hall MP, Zimmerman K, Otto P, Lubben TH, et al. NanoLuc Complementation Reporter Optimized for Accurate Measurement of Protein Interactions in Cells. 2015 [cited 17 Jul 2025]. doi:10.1021/acschembio.5b00753

6. Quijano-Rubio A, Yeh H-W, Park J, Lee H, Langan RA, Boyken SE, et al. De novo design of modular and tunable protein biosensors. Nature. 2021;591: 482–487.

7. Yeh AH-W, Norn C, Kipnis Y, Tischer D, Pellock SJ, Evans D, et al. De novo design of luciferases using deep learning. Nature. 2023;614: 774–780.

8. Yi-Hsuan Chen J, Shi Q, Peng X, de Dieu Habimana J, Wang J, Sobolewski W, et al. De novo luciferases enable multiplexed bioluminescence imaging. Chem. 2025;11. doi:10.1016/j.chempr.2024.10.013

9. Lin H, Riching K, Lai MP, Lu D, Cheng R, Qi X, et al. Lysineless HiBiT and NanoLuc Tagging Systems as Alternative Tools for Monitoring Targeted Protein Degradation. ACS Medicinal Chemistry Letters. 2024 [cited 17 Jul 2025]. doi:10.1021/acsmedchemlett.4c00271

10. LuxSitTM Pro. In: Monod Bio - Revolutionizing life sciences tools and clinical diagnostics [Internet]. Monod Bio; 21 Jul 2024 [cited 17 Jul 2025]. Available: https://monod.bio/access-luxsit-pro/

11. Dauparas J, Anishchenko I, Bennett N, Bai H, Ragotte RJ, Milles LF, et al. Robust deep learning-based protein sequence design using ProteinMPNN. Science. 2022;378: 49–56.

12. Chevalier A, Silva D-A, Rocklin GJ, Hicks DR, Vergara R, Murapa P, et al. Massively parallel de novo protein design for targeted therapeutics. Nature. 2017;550: 74–79.

